# Survival status and predictors of mortality among Breast Cancer patients in Adult Oncology Unit at Black Lion Specialized Hospital, Addis Ababa, Ethiopia, 2018

**DOI:** 10.1101/636431

**Authors:** Habtamu Abera Areri, Wondimeneh Shibabaw, Tefera Mulugeta, Yared Asmare, Tadesse Yirga

**Author notes:** Address Corresponding author, P. O. Box 4412, Addis Ababa University, Ethiopia. **Co-authors emails.** WS TM TY YA.

## Abstract

**Introduction:** Breast cancer is a foremost cause of death worldwide, ranks fifth among causes of death from all types of cancers; this is the most common cause of cancer death in women among both developing and developed countries. Breast cancer ranks first among most frequent cancers in women of Ethiopia. In spite of the high incidence, mortality rate, and survival status among breast cancer patients was not determined in Ethiopia.

**Objective:** The main aim of the study is to assess the survival status and predictor the mortality among Breast Cancer patients in Adult Oncology Unit at Black Lion Specialized Hospital in 2018.

**Methods:** An institution based retrospective follow up study was conducted in Adult Oncology Unit at Black Lion Specialized Hospital. All cases of breast cancer registered from 1^st^ January 2012 to 31^th^ December,2014 were followed for the six-year survival (until 31^th^ December, 2017). Kaplan-Meier survival curve together with log rank test was deployed to test for variations in the survival among predictor variables. Cox regression was used at 5% level of significance to determine the net effect of each independent variable on time to death of breast cancer clients.

**Results:** The results indicate that the incidence rate of mortality was 9.8 per 100 person/ years (95% CI: 8.49-11.47).The overall median survival time was 56.5(95% CI (53.46 - 60.83)) months. The overall estimated survival rate was recorded 27% (95% CI, 17.09 to 36.67 %) at 72 months of follow up, whereas at odd years (1, 3, and 5 years) were, 97.2%, 80.8%, and 46.2% respectively. Predictors of mortality were assessed at clinical stage (III&IV),(AHR =1.86), poorly differentiated histology (AHR: 3.1) & positive lymph node status (AHR:3.13),Whereas adjuvant hormone therapy (AHR: 0.67) and chemotherapy (AHR:0.72) were protective.

**Conclusion:** The overall probability of survival in Ethiopia was inferior when compared with other high and middle-income countries. Predictors of mortality were at advanced clinical stage, poorly differentiated histology grade, surgical margin involvement and positive lymph node status. In contrary, adjuvant hormone therapy, modified radical mastectomy and chemotherapy were protective factors. Hence, special emphasis could be given to early screening, stage diagnosis and initiation of treatment.

## Introduction

Breast cancer is an emerging public health danger in the worldwide, responsible for the fifth cause of death among all forms of cancers. Globally breast cancer is the second most common cancer next to lung cancer (1). Similarly, which accounts for 25% of cancer cases and 15% of cancer deaths among women (2). According to global burden of cancer 2018 estimation, globally there will be about 2.1 million new female breast cancer cases (3).

Based on 2019 estimation in U.S, an estimated 268,600 new cases of invasive breast cancer are anticipate to be diagnosed in women and about 41,760 women are likely to die(4). Globally, each year above 1.1 million women were newly diagnosed and responsible for 1.6% of female deaths from cancer causes (5). In low-income countries, breast cancer incidence was comparably low, but the most common leading cause of death among women, whereas now a day in high income countries, mortality became reduced (6). The mortality rates of breast cancer developed regions was less than the incidence because of early stage diagnosis and advanced treatment (7). Even though breast cancer has higher incidence in developed countries. But more than half of new breast cancer diagnosis and about 60% of breast cancer mortality happen in the developing countries(8).

Currently breast cancer is a rising public health problem for Sub**-S**aharan Africa (9). Likewise, in Ethiopia breast cancer became the public health disaster and one study indicate that breast cancer responsible for about 20.8 % of all cancers (10). In addition, Addis Ababa city cancer registry reports, shows that breast cancer accounts for about 34% of all female cancer cases (11). Worldwide, the cumulative 5-year survival rates of breast cancer were varying greatly, which is more than 85% in the high-income countries, while it is 60% or lower in many LMICs (12). Overall, in high-income countries, because of screening program, and early stage diagnosis, the prognosis is good. However, in low and middle-income countries, women diagnosis at advanced stage with more aggressive histological features, and related with a poorer survival (13,14).

In Sub-Sahara Africa, the 5-year survival rate of women who had breast cancer revealed that below 50%, in contrasting with 73% black women in US (9). According to GLOBOCAN data in 2012, the mortality/incidence ratios was 0.55 in Central Africa and 0.16 in the U.S (15). As a result, now a day mortality rates became escalating in certain LMICs, while it decline in most high-income countries (16). Furthermore, clients with late-stage diagnosis, limited access to standard treatment and those who present with triple-negative subtype tumors had lower survival rates (17).

In Ethiopia according to World Health Organization estimation, an age standardize incidence case of breast cancer was 12,956 and mortality rate was 25 per 100,000 women (18). As a result, to avert this burden currently the Ethiopian Federal Ministry of Health (EFMOH) prepared a task force to address the issue of non-communicable diseases with special emphasis on cancer. Overall, the fundamentals parts of the strategic framework is to reduce the incidence, mortality and improve the quality of life of cancer patients (19). Regardless of the adequate data on the incidence and survival rates of breast cancer in the western countries, survival data were scanty in Africa, and Asia countries (20-22).

In Ethiopia, most of previous studies were focused on metastasis free survival (23), trends of breast cancer (24), treatment outcome (25) and distribution of molecular subtypes of breast cancer clients (26). Despite, the government concern, to decrease the incidence and mortality rate of breast cancer patients, the survival status were still not known in Ethiopia particularly at BLSH. In line with this strategy, the current study aimed to assess the survival status and predictor of mortality among breast cancer patients at Black Lion specialized hospital, Addis Ababa, Ethiopia. Findings of this study could help to provide evidence based information to health care authorities, other stakeholders in order to design interventions that can decrease the incidence and mortality of breast cancer clients.

## Methods

### Study design, setting and population

A six-year institution based, retrospective follow up study was conducted at BLSH. Patients who have been newly diagnosed and enrolled in breast cancer treatment, from 1^st^ January, 2012 to 31^st^ December 2014 were followed until the end of the study (31^st^ December 2017). Black Lion Specialized Hospital, is located in Addis Ababa, the capital of Ethiopia. It is a teaching; central tertiary comprehensive referral hospital and has 800 beds, renders diagnostic and treatment service for about 370,000 - 400,000 patients per year. The BLSH is the only specialized hospital in the radiotherapy treatment of cancer in Ethiopia.

It has a number of services specialized in the treatment of cancers such as radiotherapy, medical oncology, anatomic pathology, nuclear medicine, gynaecology and surgery. In BLSH oncology unit, there are three senior oncologists, one palliative care specialist, nine residents, five radiotherapist, four medical Physicist, and twenty-three nurses. The most common cancer cases seen in this hospital were breast, cervical, colon and sarcomas. This study was conducted at the adult oncology unit, one of the specialty units of the hospital (27).

### Inclusion and exclusion criteria

All adult breast cancer patients who were newly diagnosed and enrolled at BLSH during the study time (*i*.*e*. 1^st^ January 2012 to 31^st^ December 2014) were included. On the other hand, women whose medical charts were incomplete, not found, those had previous diagnosis of breast cancer, a patient who had diagnosis at other hospitals and subsequently referred to BLSH for further treatment were excluded.

### Sample size and sampling technique

All breast cancer patients who attended the oncology unit of BLSH between 1^st^ January 2012 to 31^th^ December,2014 and fulfilled the inclusion criteria of the study were included in the study. This period was selected in order to have the nearest six year follow up period. A total of 1,083 breast cancer patients were registered in this period. Out of them, 627 participants who fulfilled the inclusion criteria were included in the study (**see figure1**).

**Figure 1:**
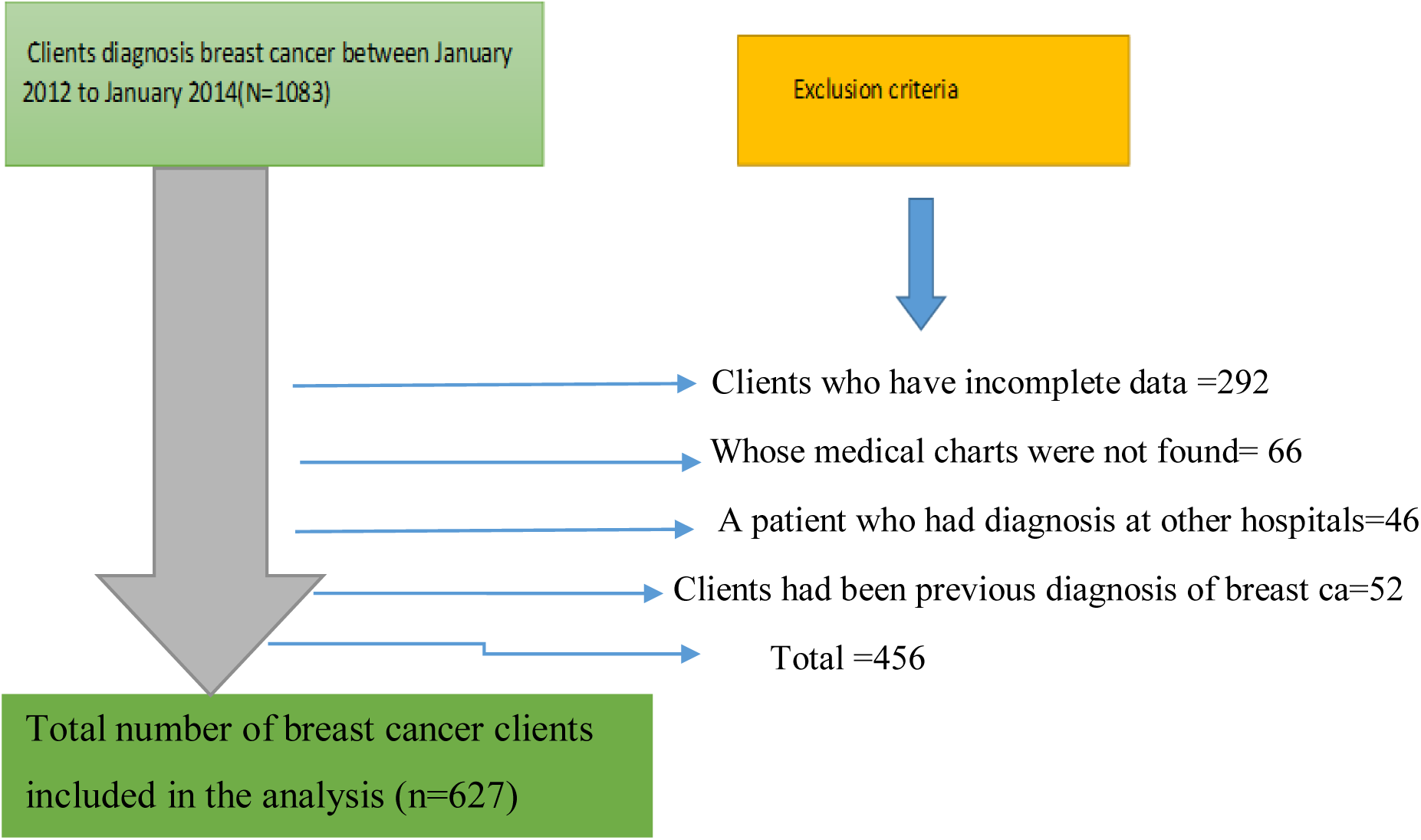
Flow diagram showing the final sample size included in the current study at black lion specialized hospital, Addis Ababa, Ethiopia, 2018.

### Data collection tool and procedure

Data collection tool were developed from different literature (5,11,16). All available information on patient records were checked and formats from different literatures were reviewed with modification. Then, appropriate data extraction format were developed in English, to extract all the relevant variables from patient charts to meet the study objectives. Training on record review was given to data collectors and supervisors for 01 day before actual data collection task on the already existing records half-day theoretical and half-day practical training. The starting point for follow-up was the time between 1^st^ January 2012 to 31^th^ December 2014 with first confirmed diagnosis of breast cancer and the endpoint was date of death, date of lost to follow up, date of last contact until 31^th^ December 2017, whichever comes first.

All sociodemographic, clinical, pathological and treatment related data were obtained from medical charts of all breast cancer cases diagnosed from 1^st^ January 2012 to 31^th^ December 2014 at black lion specialized hospital were reviewed from cancer registries. The records of all study participants were selected according to the eligibility criteria. The survival status of patients was obtained from the medical record. The time of survival was calculated as the time between the date of diagnosis of breast cancer to the date of death, or the end of study. Before collecting the data, the records were reviewed (both baseline and follow up records), death certificate complemented by registration was identified from their medical record number. The data were collected by three oncology nurses who were working at the cancer treatment center. Furthermore, the data were extracted and reviewed from the charts.

### Definition and measurement of variables

Age at diagnosis was classified into four groups (<40 years, 40-49 years, 50-59 years, >60+ years) based on the American Cancer Society facts and figures for breast cancer. Comorbidity conditions was taken from the Charlson index (28, 29). The presence of any of these conditions at diagnosis was designated as “yes”, while the absence of these condition at diagnosis was denoted as “no”. Menopausal status (Premenopausal/Postmenopausal), Marriage status (Single, Married, Widowed, Divorced), and place of residence (Urban/Rural) were also recorded.

Clinical stage at diagnosis was assigned to each patient by using the TNM classification scheme (30). In this research, the coding for those diagnosed at stages I and IV remained. Stages IIA and IIB were merged as stage II and similarly stages IIIA, IIIB and IIIC were combined as stage III. The Nottingham Grading System as grade 1, grade 2 and grade 3 measured histological grade. Tumor size was classified based on AJCC guidelines (≤2 cm, >2–5 cm, and >5cm), margins (clear/involved), axillary node status (positive/negative), Hormonal therapy (yes/no), histology type (ductal in situ, ductal invasive, lobular in situ, and lobular Invasive) were also recorded. The type of treatment was classified as chemotherapy with surgery, chemotherapy with radiotherapy, surgery alone and surgery with radiotherapy. The time (measured in months) till the death was used for the survival analysis.

### Outcome variable

The primary outcome variable was time to death of breast cancer clients in months. Breast cancer patient’s data were followed until the date of death, failure to follow-up, transferring out, or at the end of the study. Individuals who were failure to follow-up, alive and had transferred out at the end of the study period were considered as censored. The time of survival was calculated in months using the time between the dates of diagnosis to the date of the death or date of censoring.

### Data entry and analysis

Data were coded, cleaned, edited and then entered using Epi-data VS 3.1 and exported to Stata VS 14 statistical software for further analysis. Data exploration were carry out to see if there are odd codes or items that were not logical and then subsequent editing was made. Descriptive statistics were carried out in terms of central tendency for continuous data and frequency distribution for categorical data. In the current study incidence, density rate of mortality among breast cancer clients was calculated. Subsequently, the number of mortality within the follow up was divided by the total person time at risk on follow up and reported per 100 P/Y. Time to death of breast cancer clients were estimated using Kaplan-Meier survival curve and the presence of variations in survival among categories of co-variates were deployed using log rank test. Before running the cox proportional hazard regression analysis, assumption of proportional-hazard and multi-co linearity were checked.

Those independent variables, at bivariate cox regression having p-value of ≤ 0.25 was used as a cut of point to transfer in the multivariate cox regression analysis. Cox regression at 5% level of significance was used to determine the association between each independent variable to the outcome variable. The assumption of cox-proportional hazard model was checked using schoenfeld residual test and variables having P-value > 0.1 were considered as fulfilling the assumption. Hence, all the variables were fulfilling the assumption. Furthermore, residuals were checked using goodness-of-fit test with cox snell residuals, and which satisfied the model test.

## Results

### Socio-demographic characteristics

Between 1^st^ January 2012 to 31^st^ December 2014, 1,083 breast cancer patients were enrolled to Black Lion Specialized Hospital, of them, 627 were eligible for this study. Among the study participants, 458 were censored and the rest 169 develop the event. The mean age of study participants at the time of breast cancer diagnosis was 42.61 years with SD ± 12.28 years. Almost nearly half, 279 (44.5%) of the age group was less than 40 years old. About two-thirds, 403 (64.3%) of patients were married; most, 433 (69.1%) of the women was premenopausal (age less than 50 years old); slightly more than one-third, 224 (35.7%) were having preexisting medical problem during diagnosis. More than half, 366 (58.4%) of the participants were urbanites are shown below (**Table 1**).

**Table 1:** Socio-demographic characteristics of breast cancer patients at Black Lion Specialized Hospital, Addis Ababa, Ethiopia, from 1^st^ January 2012 to 31^st^ December 2017 (n=627).

| Covariate | category | status at last contact |  | Total<br>No. (%) |
| --- | --- | --- | --- | --- |
|  |  | Censored<br>No. (%) | Death<br>No. (%) |  |
| Age in years at time of diagnosis | <40 | 212(75.9) | 67(24.1) | 279(44.49) |
|  | 40-49 | 111(68.09) | 52(31.91) | 163(26.0) |
|  | 50-59 | 93(78.81) | 25(21.19) | 118(18.82) |
|  | >60+ | 42(62.28) | 25(37.31) | 67(10.68) |
| Place of residence | Urban | 285(77.86) | 81(22.14) | 366(58.37) |
|  | Rural | 173(66.28) | 88(33.71) | 261(41.63) |
| Marital status | Single | 71(76.34) | 22(23.66) | 93(4.83) |
|  | Married | 301(74.68) | 102(25.32) | 403(64.27) |
|  | Divorced | 63(63.64) | 36(36.36) | 99(15.78) |
|  | Widowed | 23(71.87) | 9(28.13) | 32(5.1) |
| Menopause status | Premenopausal | 316(72.97) | 117(27.03) | 433(69.05) |
|  | Postmenopausal | 142(73.19) | 52(26.81) | 194(30.95) |
| Preexisting medical diagnosis | No | 323(80.14) | 80(19.86) | 403(64.27) |
|  | Yes | 135(60.26) | 89(39.74) | 224(35.73) |

### Clinical, Histopathological and treatment characteristics

More than half, 357 (57%) of the women were clinical stage III at the time of diagnosis. Invasive Ductal Carcinoma (IDC) was predominant, 427 (68.1%) histology type of cancer. Nearly half, 304 (48.5%) of the histology grade were moderately differentiated; Surgery associated with radiotherapy was the common mode of treatment which accounts for 256 (40.8%). More than half, 369 (58.85%) of tumor size was less than 2.5 cm on presentation. About two third, 407 (64.91 %) of the study participants were reported receiving hormone therapy. Nearly half, 229 (48.11%) of patients had modified radical mastectomy. The clinical, histopathological and treatment characteristics of the study participants are shown below (**Table 2**).

**Table 2:** Baseline clinical, histological and treatment information of breast cancer patients at Black Lion Hospital, Addis Ababa, Ethiopia, from 1^st^ January 2012 to 31^st^ December 2017 (n=627)

| Variable | Category | Vital status at last contact |  | Total<br>No. (%) |
| --- | --- | --- | --- | --- |
|  |  | Censored<br>No. (%) | Death<br>No. (%) |  |
| Stage of breast cancer | I | 21(95.45) | 1 | 22(3.5) |
|  | II | 145(86.82) | 22(13.18) | 167(26.63) |
|  | III | 245(68.62) | 112(31.38) | 357(56.93) |
|  | IV | 47(58.02) | 34(41.98) | 81(12.91) |
| Histology grade | Grade I | 84(91.3) | 8 | 92(14.67) |
|  | Grade II | 237(77.96) | 67(22.04) | 304(48.48) |
|  | Grade III | 137(59.3) | 94(40.7) | 231(36.84) |
| Histology type | Ductal insitu | 136(83.44) | 27(16.56) | 163(26.3) |
|  | Invasive ductal | 294(68.85) | 133(31.15) | 427(68.1) |
|  | Lobular insitu | 15(75) | 5 | 20(3.2) |
|  | Invasive lobular | 11(73.4) | 4 | 15(2.4) |
| Deep surgical margin | Free | 393(80.2) | 97(19.8) | 490(79.28) |
|  | involved | 58(45.3) | 70(54.7) | 128(20.72) |
| Node status | Negative | 364(80) | 91(20) | 455(72.68) |
|  | Positive | 93(54.38) | 78(45.62) | 171(27.32) |
| Metastasis at time diagnosis | No | 348(87.21) | 51(12.79) | 399(63.64) |
|  | Yes | 110(48.24) | 118(51.76) | 228(36.36) |
| Tumor size | ≤2.5cm | 289 (81.17) | 67(18.82) | 356(56.78) |
|  | 2.5-5cm | 152(63.34) | 88(36.66) | 240(38.27) |
|  | >5cm | 17(54.8) | 14(45.2) | 31 (4.95) |
| Number of Positive node | <2 | 284(75.53) | 92(24.47) | 376(59.97) |
|  | ≥2 | 174(69.32) | 77(30.68) | 251(40.03) |
| Treatment mode | Surgery associated with radiotherapy <sup>a</sup> | 165(64.45) | 66(25.65) | 256(40.82) |
|  | Surgery<sup>b</sup> | 101(72.66) | 38(27.34) | 139(22.18) |
|  | others<sup>c</sup> | 191(82.68) | 65(28.13) | 231 (36.8) |
| Type of surgery | Breast-conserving surgery | 65 (58.56) | 46(41.44) | 111(23.32) |
|  | total mastectomy | 86 (76.1) | 27 (23.89) | 113(23.74) |
|  | Modified radical mast | 180 (78.6) | 49 (21.4) | 229(48.11) |
|  | Axillary node dissection | 14(60.86) | 9(39.13) | 23(4.83) |
| Chemotherapy | No | 67(72.8) | 25(27.2) | 92 (14.7) |
|  | Yes | 391 (73.1) | 144(26.9) | 535(85.3) |
| Endocrine therapy | No | 137(62.27) | 83(37.72) | 220(35.08) |
|  | Yes | 321(78.86) | 86(21.13) | 407(64.92) |

<sup>a</sup>Surgery associated with radiotherapy (or also combined with chemotherapy)
<sup>b</sup>Surgery only (or also combined with chemotherapy) ;<sup>c</sup> radiotherapy with chemotherapy( surgery therapy).

### Survival status of breast cancer patients

The overall mortality rate in the cohort during the 1,712 person-years of observation (PYO) was 9.87 per 100 (95% CI: 8.49-11.47) person-years follow up. The cumulative incidence of this study was 169 (27%) with the confidence interval (95% CI, 23.6-30.3%) of patients were died over six years. However, 458 (73.04%) were censored till the end of the study. Of these, 194 (42.27%) were failure to follow up, 132 (28.76%) were alive, 119 (25.9%) was against medical advice and the rest were transferred to other institution during the overall study follow up period.

### Survival probability of breast cancer patients

In the current study, 627 breast cancer participants were followed for a total of 72 months. The estimated median survival time was 56.5 (95% CI, 53.46 - 60.83) months. Kaplan-Meier survival estimation showed that overall estimated survival rate after diagnosis of breast cancer was 26.42% (95% CI, 17.09 to 36.67 %) at 72 months of follow up. Likewise, the estimated cumulative survival rate was 97.2%, 89.8%, 80.8%, 66.33%, 46.2% and 26.4% at 12, 24, 36, 48, 60 and 72 months respectively **(see figure 2)**.

**Figure 2.**
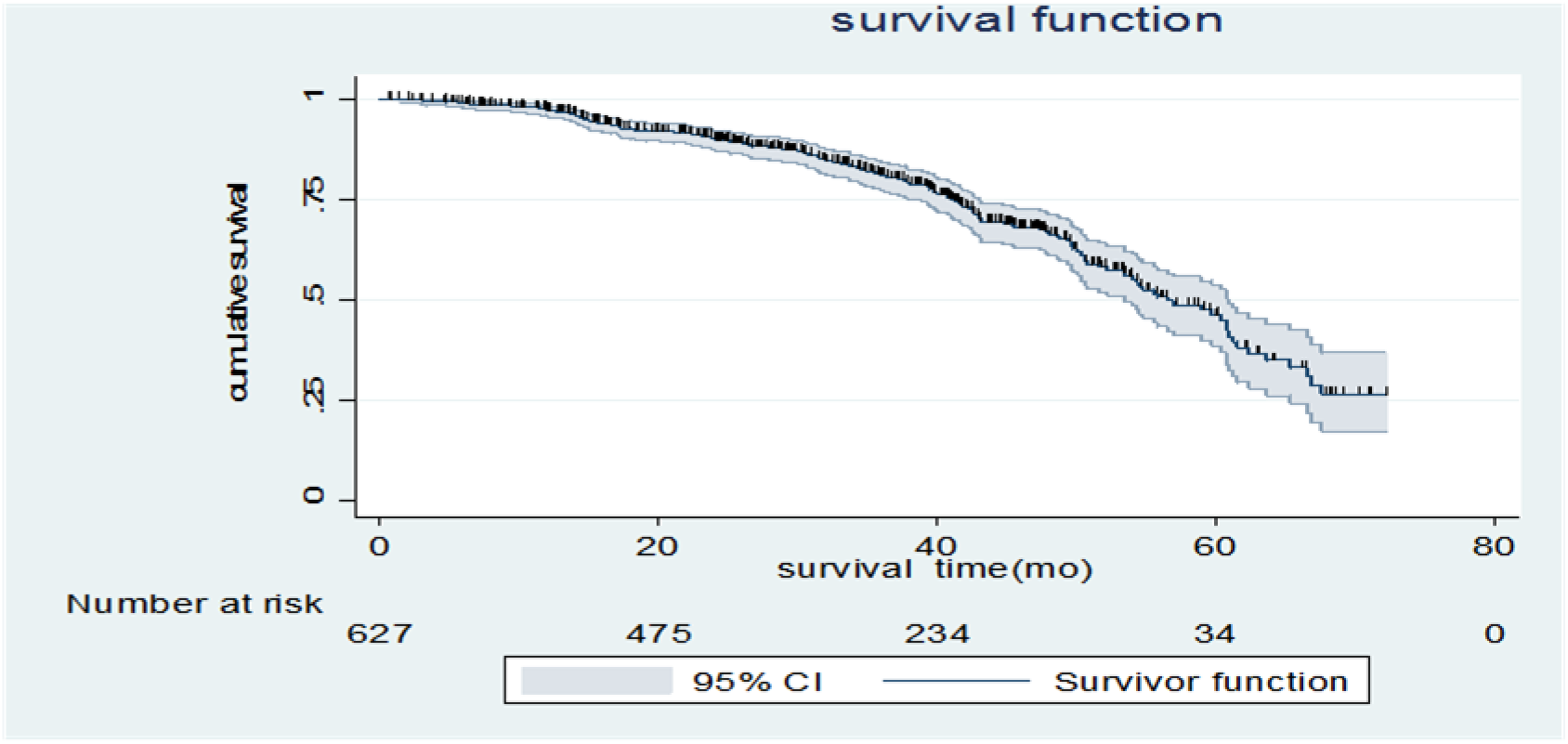
Kaplan-Meir survival estimate on the overall survival time of breast cancer clients at Black Lion Specialized Hospital, Addis Ababa, Ethiopia, 2018 (n = 627).

### Survival function among different predictors

Log-rank test was carried out to check the presence of any significant differences in survival time among various levels of the categorical variables. According to the present study it was found that the median survival time for those who had clinical stage I, II or III at baseline had a longer survival time than those who had advanced clinical stage (IV) (45.6 months, 95% :41.04-50.57) at baseline with a p-value < 0.000. The median survival time for those who have negative lymph node status had a longer survival time 63.6 months than those who had positive lymph node at baseline (47.34 months, 95% CI: 41.2-49.7),which is supported by the p-value of 0.000.

Among cases diagnosed in early stage (I & II), cumulative survival was 56.65 % (95 % CI: 35.94-72.94 %), while those cases detected at late stage (III&IV), had a survival rate of 18.49% (95 % CI: 10.03-28.98%) with the testing equality among the groups with p value of 0.000. A 6-year survival rate for those diagnosed with IDC (invasive ductal carcinoma) was 20.2% (95%CI,10.82-30%) which is lower as compared to other histological types 43.26%(22.39-62.56) shown below (**table 3 and figure 3**).

**Table 3:** Survival time, cumulative survival probability, significance and log rank test for the study population according to different characteristics of patients during 6-year of follow-up (Kaplan-Meier Method) of breast cancer clients at Black Lion Specialized Hospital, Ethiopia from 1^st^ January 2012 to 31^st^ December 2017 (n=627).

| Covariate | Survival time, mo, (95% CI) | Overall 6-year survival probability (%) | Log rank test (p-value) |
| --- | --- | --- | --- |
|  | Median(95%CI) |  |  |
| Residence |  |  |  |
| Urban | 59.7 (54.47-..) | 36.57 | (8.83)* |
| Rural | 50.73 (49.3 -60.67) | 17.26 |  |
| Menopause status |  |  |  |
| Premenopausal | 60.7 (53.47- 63.63) | 28.56 | 0.63 |
| Postmenopausal | 54.5 (48.64-58.84) | 19.67 |  |
| comorbidity |  |  |  |
| No | 63.64 (60.84- .) | 35.62 | (23.18)* |
| Yes | 50.04 (45.6-53.37) | 12.89 |  |
| Stage |  |  |  |
| I |  | 92.86 | (41.46)* |
| II | 61.54 (59.7-..) | 43.56 |  |
| III | 54.46 (50.24-60.67) | 20.18 |  |
| IV | 45.6 (41.04- 50.57) | 16.8 |  |
| histology grade |  |  |  |
| Grade I |  | 66.4 | (62.38)* |
| Grade II | 60.74 (55.76- 67.54) | 34.6 |  |
| Grade III | 48.63 (42.96 -50.73) | 0 |  |
| Deep surgical margin |  |  |  |
| free |  |  | (56.32)* |
| involved | 62.3 (59.7- .) | 41.31 |  |
|  | 42.97 (39.74-48.34) | 2.93 |  |
| Number of involved lymph nodes |  |  |  |
| <2 | 60.74 (54.47-66.5) | 30.21 | 3.65 |
| >=2 | 53.96 (49.6-59.7) | 18.93 |  |
| Tumor size |  |  |  |
| ≤2.5 | 61.53 (60.67- ...) | 45.65 | (26.42)* |
| 2-5 | 50.57 (47.96-54.33) | 11.35 |  |
| >5 | 45.36 (41.84-...) | 0 |  |
| Metastasis at diagnosis |  |  |  |
| No | 66.5 (60.84- .) | 48.5 | (99.9)* |
| Yes | 42.5 (40.04- 47.34) | 5.4 |  |
| Endocrine therapy |  |  |  |
| No | 47.97 (41.84-51.64) | 6.8 | (40.65)* |
| Yes | 65.3 (58.84- .) | 40.58 |  |
| Lymph node status |  |  |  |
| Negative | 63.64 (60.67- .) | 38.27 | (48.95)* |
| Positive | 47.34 (41.2-49.7) | 5.1 |  |
| Neo adjuvant |  |  |  |
| Chemotherapy |  |  |  |
| No | 53.47 (42.5- | 24.9 | 2.35 |
| Yes | 58.84 (53.97-61.54 | 27.52 |  |
| Type of surgery |  |  |  |
| Breast conserving |  |  |  |
| surgery | 49.6 (43.03-54.34) | 12.6 | (19.3)* |
| Total mastectomy | 60.74 (51.64- .) | 39.2 |  |
| MRM | 62.3 (58.84- .) | 39.4 |  |
| Axillary dissection | 50.03 (14.64- .) | 0 |  |

mo; months, CI; confidence interval, SE: Standard Error, MRM; Modified Radical Mastectomy,
\* indicates that the variables have significantly difference in survival among groups at 95% level of significant ( $P < 0.05$ ).

**Table 4:**
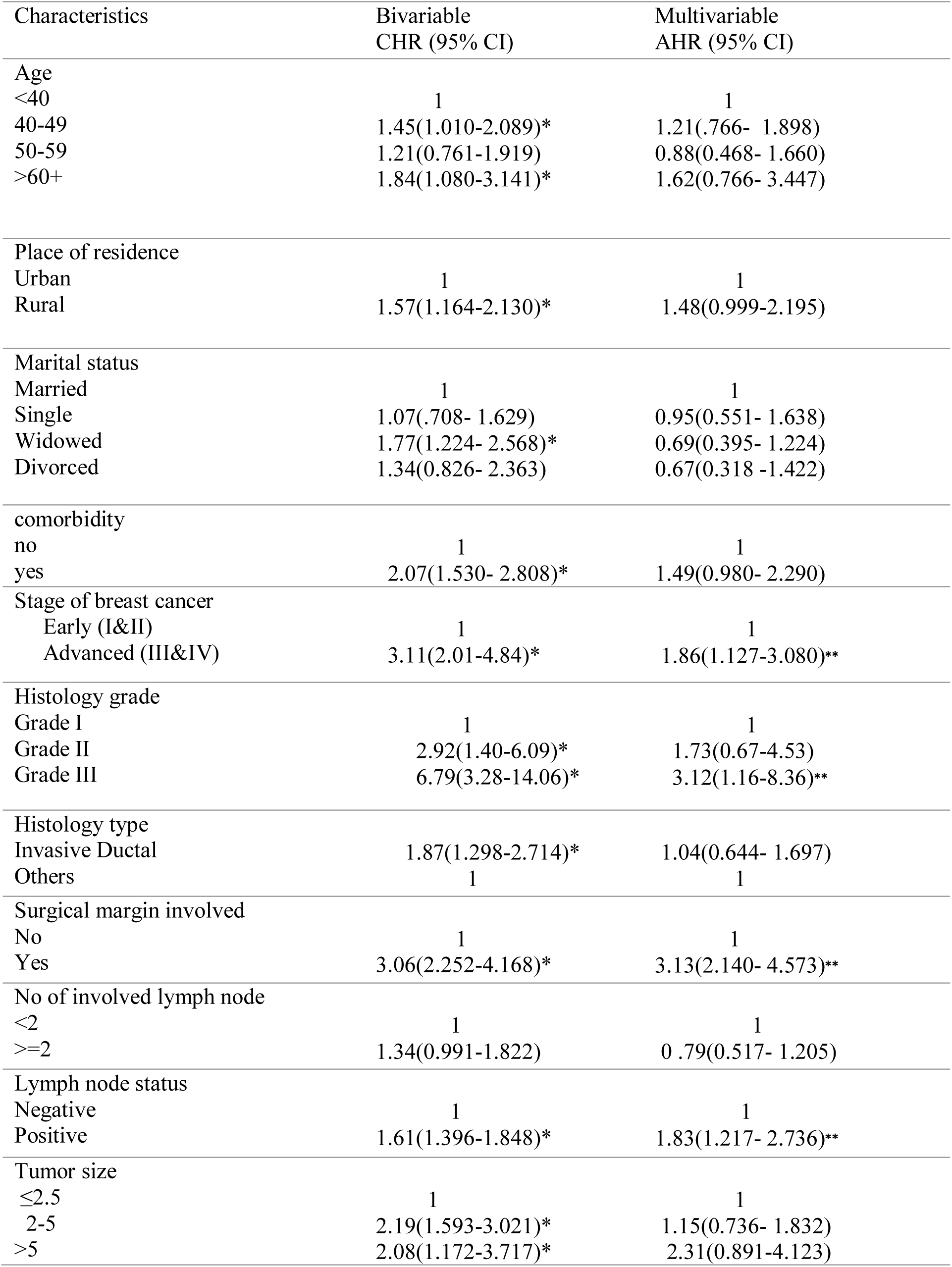
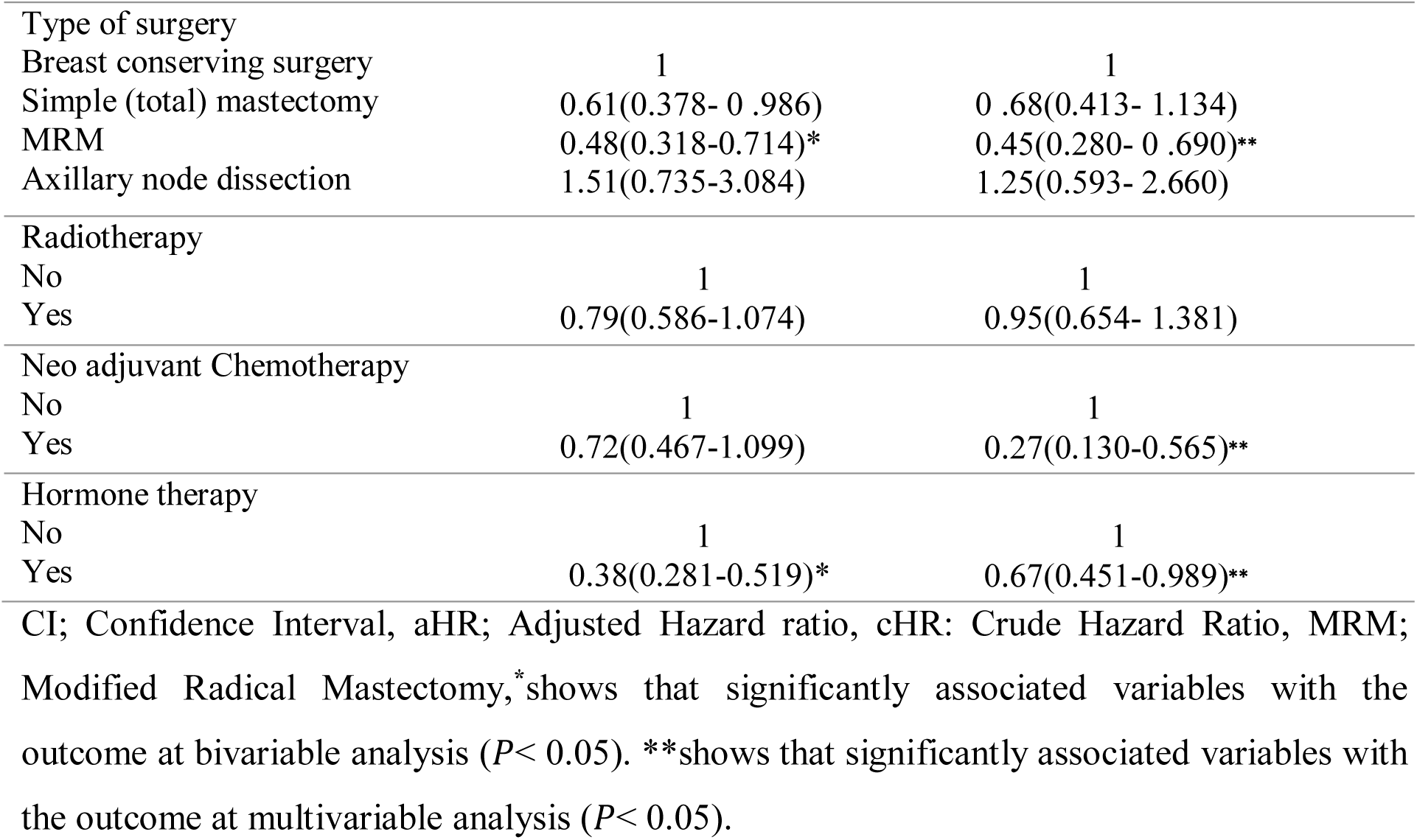
Results shows cox proportional hazard regression analysis to identify predictors of mortality among breast cancer patients at Black Lion Specialized Hospital, Addis Ababa, Ethiopia, 1^st^ January 2012 to 31^st^ December 2017 (n=627).

**Figure 3.**
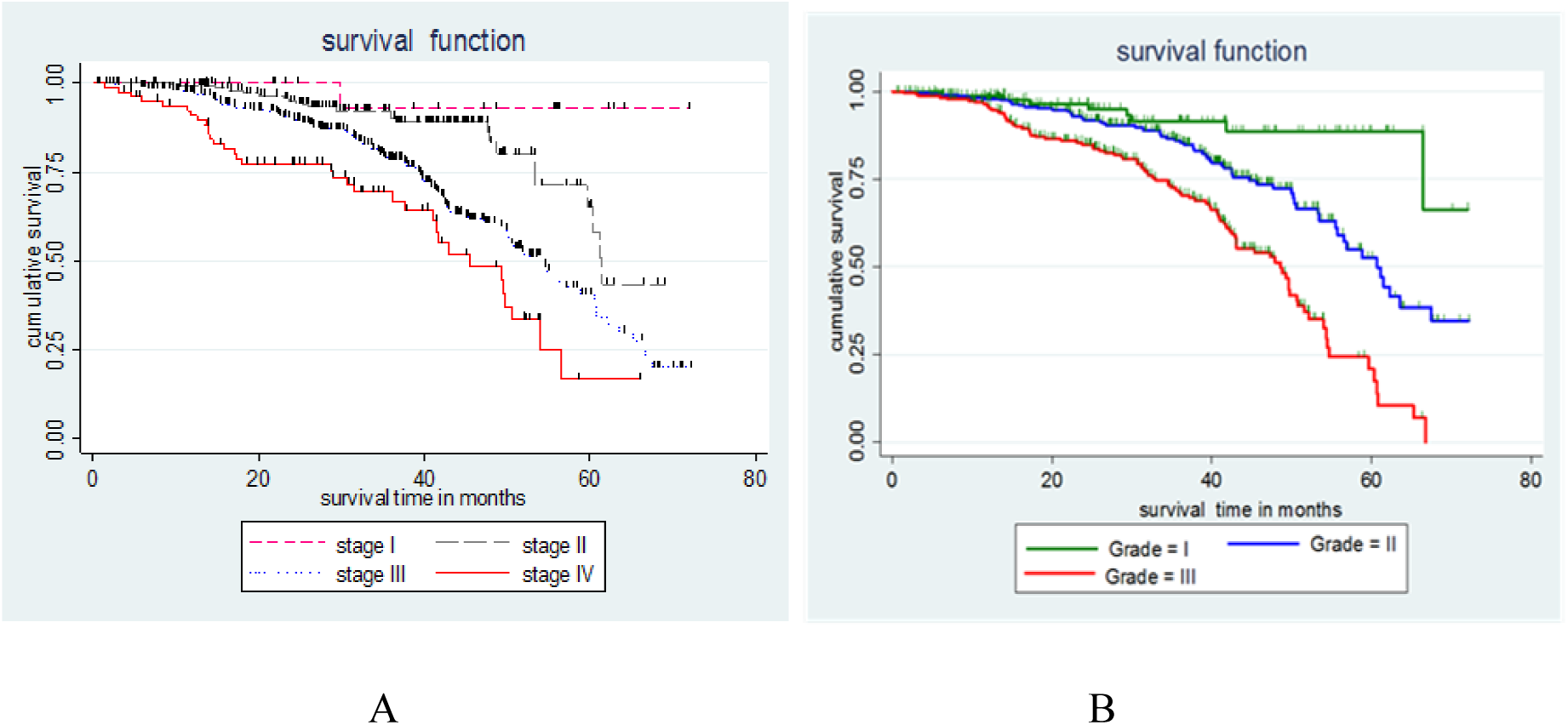
Kaplan-Meier Survival function among different groups of breast cancer clients (A), stage at diagnosis (B), histologic grade at diagnosis at Black Lion Specialized Hospital, Addis Ababa, Ethiopia, 2018 (n=627).

### Testing overall model fit

In order to check the overall model fit we compare the jagged line with the reference line, and we observe that, the cox model fit these data to reasonable extent. In addition, if the cox regression model fits the data, these residuals should have a standard censored exponential distribution with hazard ratio. Overall, the hazard function follows the 45° line very closely. Hence, the output shows cox–snell residuals were satisfied the overall model fitness test (**see figure 4**).

**Figure 4.**
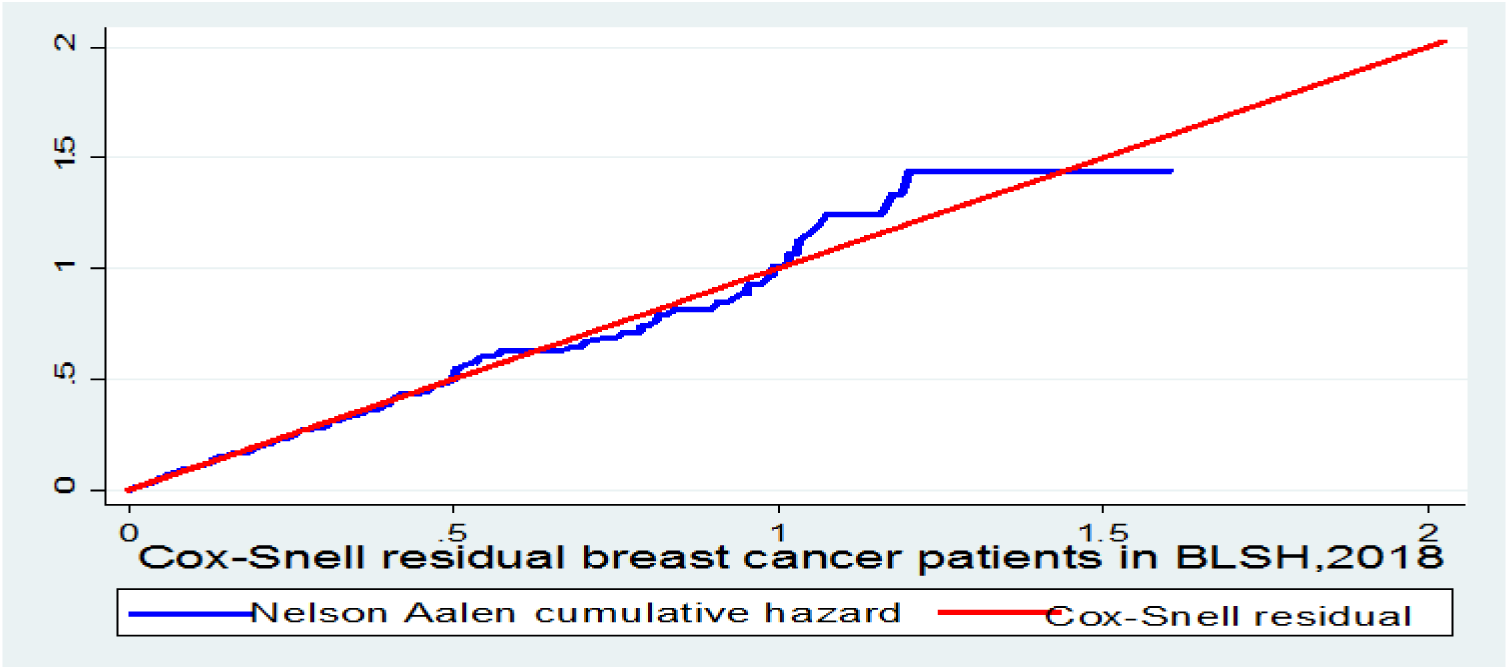
This figure shows cox-snell residual Nelson -Aalen cumulative hazard graph on breast cancer clients at Black Lion Specialized Hospital, Addis Ababa, Ethiopia, 2018.

### Predictors of mortality

In multivariable cox regression analysis explanatory variables with p-value < 0.25 at the bivariate analysis and non-collinear independent variables were included. The result of multivariable analysis revealed that women with late stage of breast cancer (III and IV) were 1.86 times higher hazard of death than those who had early stage breast cancer at the time of diagnosis (I and II) (AHR : 1.86, 95%CI: 1.13-3.08). Those women whose surgical margin were involved at baseline 3.13 times more likely to die than those women whose surgical margin was not involved (AHR: 3.13, 95% CI: 2.140-4.573).

Patients having positive lymph node status were 1.83 times higher hazard of death as compared to those having negative lymph node at time of diagnosis (AHR: 1.83, 95% CI:1.217-2.736). Women those who are being histologic grade III at the beginning of breast cancer diagnosis were 3.12 times higher risk of death as compared to those who have grade I histology(AHR: 3.12, 95% CI:1.16-8.39). Furthermore, women who have taken endocrine therapy during the six year follow up time have reduced mortality by 33% compared to those who are not taken endocrine therapy (AHR: 0.67, 95% CI: 0.451-0.989).

## Discussion

In the current study, the overall mortality rate of breast cancer patients were 9.8/100 women-years. Our finding was much higher than the study done in Brazil (31), where age-standardized mortality rate within 6-year follow up period was 431.8/100 000 women-years. The observed variation in mortality rates might be because of the fact that the more advanced stage (III & IV) breast cancer clients were presented in our finding. Other possible explanation could be methodology difference; they have been used ecological analysis.

In this 6-year retrospective follow up study, the cumulative survival probability at 1, 3, and 5 years were 97.2%, 80.8% and 46.2% respectively. This finding is in line with studies conducted in Malaysia (70.8%, 56.9% and 49.4%) (32) and Ghana, (47.9%) (21). However, this finding is higher than in Tanzania (21.8%) (33), Cameroon (30%)(34), Uganda (51.8 %) (35) and Vietnam, (94%, 83% and 74%)(5). On the contrary, survival probability is still lower when compared with studies done in Northwest Iran (96%, 86%, and 81%), Germany (83%) and the Qidong (83.61%, 67.53% and 58.75%) respectively (36-38). Similarly, the overall cumulative 6 years estimated survival rate of the current study was 26.42 % (95 % CI: 17.09-36.67 %) at 72 months of follow up. This result is inferior than the study done in Qidong (56.04%) (38).

This gap might be due to several reasons. This perhaps could be due to advanced stage at diagnosis (69.5%). However, 52.3% in Ghana, 27.6% in Vietnam, 67.8% in Cameroon is due to lack of early screening programs, limited treatment facilities, and financial problems. Other explanation could be the facilities to treat cancer are located in the capital city of the country. This could result in delay of diagnosis and treatment. Additionally, possible explanation on discrepancies in survival might be due to different methodologies applied in each studies, as well as variances in local cancer care. Furthermore, the molecular and genetic differences among breast cancer clients may also contribute to variations in survival probability.

In the current study, the overall median survival for histologically confirmed breast cancer was 56.5 (95% CI, 53.46 - 60.83) months. The results were higher than the previous studies in which the median survival was 24 months in Cameroon (34), 40 months in Sudan (39), but lower than the results reported from Malaysia the median survival was 68.1 months (32). Although our study showed higher overall median survival than that of the Cameroon and Sudan studies, the lower value obtained in comparison with Malaysia is most probably due to the shorter period of study. In addition, it could be difference in health seeking behavior and treatment adherence among those studies.

In the log rank test, it is observed that the median survival time for those who had clinical stage I, II or III at baseline had a longer survival time than those in advance clinical stage (IV) (45.6 months, 95% :41.04-50.57). Similarly, the median survival time for those who have negative lymph node status had a longer survival time (63.6month, 95%CI: 60.67-) than those who had positive lymph node at baseline (47.34 months, 95% CI: 41.2-49.7). These values are in line with the studies conducted in different African countries (39,40).

In the present study, women who had positive lymph node at diagnosis were nearly 2 times prone for death than women who had negative lymph node at baseline. These results were in agreement with different studies conducted in African countries (21,41) and the study done in Iran (42) in which lymph node status was an important determinant of survival. In addition, as the number of lymph nodes increases, also does the relapse rate, while the survival rate decreases. Women who had histologic grade III at diagnosis were nearly 3 times more likely to die as compared to women who had well differentiated histologic (grade I) at baseline. These results were in agreement with the studies conducted in different Asian countries (32, 42, 43).

According to the results obtained, women who had positive surgical margin were 3.13 times more prone to death than women who were free from surgical margin at time of diagnosis. Most of the previous studies go hand in hand with the present findings (5,35). These findings might be explained by the fact that the residual disease at the surgical margins could increase the risk of local recurrence and possibly death through the years.

As shown by others studies, women who had advanced stage of breast cancer (III and IV) at diagnosis were nearly 2 times more vulnerable to death than women who had early stage breast cancer (I&II) at time of diagnosis. The same line of results were presented by several studies conducted in different African and Asian countries (5,35, 37,43). This reflect that stage was a major predictor of mortality among breast cancer clients and had the most significant influence on patient outcome in the present study.

The results indicate that patients who had undergone modified radical mastectomy (MRM) experienced 55% reduced chances of mortality than those who underwent breast-conserving surgery. The observed results are in contrary with the studies done in different African and Asian countries (34,36,42). The difference in survival between two groups could be because of late stages at presentation and poor infrastructure for treatment of breast cancer has made this approach of surgical treatment more common in many developing countries including Ethiopia. Scientifically, breast-conserving surgery is usually recommended for early stage breast cancer in contrast to modified radical mastectomy. Other possible explanation could be the options for treating breast cancer patient, which depends on the stage of disease.

Unlike most of previous findings (40,44), those women who were receiving neo-adjuvant chemotherapy was found to be reduce the chances of death by 73% than those patients without neo-adjuvant chemotherapy. However, the current result is in agreement with study done in France (45), which revealed a 25% reduction in the relative risk of death. Overall, the present study found a protective effect of neo-adjuvant chemotherapy on survival of breast cancer patients. This might be due to patients with advanced stage breast cancer, neo-adjuvant chemotherapy would be applied in order to shrink the tumor size and facilitate surgery. In our study, the better survival rate was related to patients receiving neo-adjuvant chemotherapy. Additional studies should be conducted to confirm a protective effect of neo-adjuvant chemotherapy on survival rate of breast cancer clients.

The final predictor of survival identified in the study was hormone therapy, which reduced the chances of death by 33% than those who could not undergo hormone therapy. The findings are in consistent with previous findings that were conducted in different continents (5, 40, 44) which reflect that the effect of hormone therapy on survival rate was often stated along with the impact of hormone receptors.

This study has some limitations too. First, cause specific survival was not determined due to lack of data on specific cause of death. This may over estimate breast cancer related mortality rate. Second, limited data on hormone receptors; estrogen receptor, progesterone receptor and Human Epidermal Growth Factor Receptor 2 prevented us from analyzing the role of them on survival time. Likewise, data were not recorded on socio economic status, such as occupation and educational level. Selection bias was possibly introduced during secondary data collection because patient’s with incomplete charts were excluded. Moreover, the data were collected over the period 1^st^ January 2012 to 31^st^ December 2014 and do not reflect current utilization of advanced treatment methods and new medications for breast cancer treatment.

## Conclusion

The overall probability of survival in patient’s diagnosis of breast cancer was 27% at 72 months of follow up in Ethiopia that is inferior when compared with those of high and middle-income countries. Significant predictors of mortality after diagnosis of breast cancer were advanced clinical stage, grade III histology, involvement of surgical margin and positive lymph node status. In contrast, hormone therapy, modified radical mastectomy and chemotherapy reduced mortality. Hence, a special emphasis should be given to early screening, diagnosis and initiation of treatment before advanced stage. leading to high mortality.

## Abbreviations

AAU: Addis Ababa University,
AJCC: American Joint Committee on Cancer,
BLSH: Black Lion Specialized Hospital,
ER: Estrogen Receptor,
EFMOH: Ethiopian Federal Ministry of Health,
GLOBOCAN: Global Burden of Cancer,
HER2: Human Epidermal Growth Factor Receptor 2,
HIC: High Income Country,
IDC: Invasive Ductal Carcinoma,
LMICs: Low and Middle Income Countries,
PR: Progesterone Receptor,
SSA: Sub-Saharan Africa.

## Ethics approval and consent to participate

Ethical clearance was obtained from Addis Ababa University, ethical review board. Permission for conducting the study was gained from Adult Oncology Unit, Black Lion Specialized Hospital. The data were collected from the client charts, as a result the individual clients were not exposed to any harm as far as the privacy was kept. To keep the confidentiality, all collected data was coded and locked in a separate room before being entered into the computer.

## Consent for publication

Not applicable.

## Availability of data and materials

Data will be shared upon request and will be obtained by emailing to the corresponding author using ““.

## Authors’ contributions

WS, TM, and HA were involved in conceptualize, design, methodology, statistical analysis, developing the initial drafts of the manuscript and revising. Whereas YA and TY involved in software, supervision and subsequent drafts. In addition, WS, TM, and HA prepared the final draft of the manuscript, reviewed and edited. All authors read and agreed on the final draft of the manuscript.

## Acknowledgment

We would like to acknowledge Addis Ababa University for funding and Black Lion Specialized Hospital for permission to undertake this study. Our heartfelt thanks to Black Lion Specialized Hospital Managers, all oncology unit staffs, card room officer and data collectors for their cooperation during data collection.

